# Low Level LASER Therapy Induces Therapeutic Angiogenesis in Diabetic Mice with Hindlimb Ischemia

**DOI:** 10.1101/496208

**Authors:** Shi-Jie Huang, Yi-Hsien Teng, Yu-Jung Cheng

## Abstract

Patients with diabetes mellitus (DM) are at high risk of developing peripheral arterial obstructive disease (PAOD) in lower extremities. Previous studies show low level LASER therapy (LLLT) can increase angiogenesis *in vivo* and *in vitro*. Here we performed hindlimb ischemia as PAOD model on diabetic mice to test the effects of LLLT. Twenty C57Bl/6 mice were randomly divided into four groups. Control group mice received femoral artery ligation/excision only. DM group mice were injected with Streptozocin (STZ) to induce diabetes followed by femoral artery ligation/excision. LLLT group mice received femoral artery ligation/excision and the lower limbs received LASER treatment for five days (660nm, 10 min, 1.91 J/cm^2^) started from second day postoperatively. DM+LLLT group mice were received femoral artery ligation/excision and LASER treatment after diabetes induced. Three days after LASER treatment finished, limb blood flow was measured by Laser Doppler perfusion imaging. Capillary density was assessed by immunofluorescence staining. CD31, VEGF, HIF-1α, phospho-ERK, iNOS and eNOS protein level was examined by Western blot. Blood perfusion, capillary density, CD31, and VEGF protein levels were significantly higher in those groups received LLLT compared to control and DM group. Low level LASER significantly increased ERK phosphorylation and HIF-1α expression. In addition, phospho-eNOS was increased but iNOS protein level was decreased in mice received LASER treatment. In summary, the ability of low level LASER to induce therapeutic angiogenesis in diabetic mice suggested this approach deserves investigation as a novel approach to treat PAOD patients.

## Introduction

Lower limb ischemia is often found in patients with diabetes mellitus (DM), and peripheral arterial obstructive disease (PAOD) is one of major health care problems. The age-adjusted prevalence of PAOD is 12%, and the prevalence of PAOD can be expected to increase because of the aging population[1]. The primary pathophysiology of PAOD is arterial stenosis or occlusion leading impair perfusion in lower extremities. Symptomatic PAOD may cause intermittent claudication, which induces pain during movements in affected limb, especially ambulation. In advanced stage of PAOD, critical limb ischemia cause pain at rest and ulcer formation. These different stages of PAOD produce functional impairments and restricts patient’s activity of daily livings.

The treatments for critical PAOD, which target improving longer-term cardiovascular outcomes, include risk modification, exercise, wound management, endovascular and surgical interventions[2]. Exercise can increase maximum walking time, improve walking ability, and gait patterns in non-surgical patients[3]. Antiplatelet and lipid lowering therapy delay the rate of PAD progression and reduce the incidence of new onset intermittent claudication[4, 5]. In patients with critical limb ischemia, therapeutic interventions such as revascularization and surgery have to be considered. Endovascular treatment using percutaneous transluminal angioplasty can lower morbidity and mortality in patients with femoropopliteal artery stenosis [6]. The aims of surgical interventions are preventing limb loss, improving ulcer healing, activity of daily living, quality of life, and survival rate[6]. In addition, therapeutic angiogenesis was investigated whether it can ameliorate pain and improve exercise capacity in patients with critical PAOD [7].

Angiogenesis is the growth and development of new capillaries from a pre-exiting vasculature, and therapeutic angiogenesis seeks to harness this phenomenon for the treatment of disorders of inadequate tissue perfusion[8]. LLLT, which is wildly used in the physical therapy field, is the application of light in the range of 1mW-500mW. Physical therapists use LLLT to promote wound healing, reduce inflammation, and relieve pain [9–12]. Unlike other medical laser procedures, LLLT does not act via thermal mechanisms but rather through a photochemical effect. Evidence has shown that LLLT can alter cell and tissue function, including stimulating collagen production [13], promoting DNA synthesis [14], and increasing ATP content [15]. On endothelial cells, LLLT stimulates cell proliferation, vascular endothelial growth factor (VEGF) expression, and angiogenesis [16, 17]. In our previous study, we found LLLT can prevent endothelia cells from TNF-α-induced apoptosis. Due to damage of vascular endothelium is the major consequence of exposure to inflammatory cytokines, LLLT might improve wound healing via preventing endothelial cells from inflammation-induced apoptosis[18].

Here we demonstrated that this LLLT could increase the expression of VEGF, CD31, and capillary density to hindlimb ischemia in a mouse model of PAOD. We therefore further evaluating the effects of Low level LASER on angiogenic and mitogenic related signals.

## Material and methods

### Reagents

Streptozotocin was purchased from Tocris (Tocris Bioscience, Ellisville, MO); anti-eNOS, anti-phospho-eNOS antibodies were obtained from BD Bioscience (San Diego, USA); anti-phospho-ERK, anti-ERK antibodies were purchased from Cell Signaling Technology (Beverly, MA). HIF-1α antibody was obtained from Novus (Novus Biologicals, Littleton, CO). CD-31 endothelial cell antibody was purchased from eBioscience (eBioscience, San Diego, USA) and collagen type IV antibody was from Abcam (Abcam, Cambridge, MA, USA).

### Diabetes induction and hindlimb ischemia model

Eight weeks old C57Bl6 mice weighing 25-30g were housed under 12 h light/dark cycle at 23 ±2 °C. Diabetes was induced by intraperitoneal injection of STZ (50 mg/kg, dissolved in 0.05 M cold sodium citrate buffer, pH 4.5) for five consecutive days. The normal control animals received only citrate buffer. One month later, the diabetic animals (blood glucose level ≥200 mg/dl) and control mice were then received hindlimb ischemia surgery (HLI). After anesthesia, an incision was made along the left medial thigh to allow isolation, ligation, and excision of the femoral artery from its origin just above the inguinal ligament to its bifurcation at the origin of the saphenous and popliteal arteries[19]. After wound closed, the perfusion of hindlimb was measured by LASER Doppler (moorLDI2-IR, Moor Instruments, Devon, UK).

After 3 days of ischemia, the mice were anesthetized with isoflurane. Lower limbs were exposed to AlGaInP-diode laser (AM-800, Konftec Co., Taipei, Taiwan) for five days. The probe of the laser device was fixed vertically, 30 cm above the mice. Laser irradiation was applied with a wavelength of 660 nm at an output power of 50 mW/cm^2^ for 10 minutes. The average energy density each day of the treatment was 1.91 J/cm^2^.

### Tissue preparation and histological section

Before tissue harvest, bilateral hindlimb perfusion measures were performed after anesthesia with isoflurane. After mice were sacrificed, the ischemic and contralateral gastrocnemius/soleus muscles were harvested from tendon to tendon. Half muscle tissue was embedded in O.T.C and quickly freeze with liquid nitrogen. The tissue was processed to frozen cryostat sections (7 μm). The other half tissue was collected and stored at −80°C for further protein analysis.

### Capillary density and immunofluorescence staining

Capillary density (capillaries/mm^2^) was measured by counting five random fields (×100 magnification) from CD31/Collagen IV double staining [20]. Immunofluorescence staining images were captured and analyzed using fluorescence Olympus BX43 microscopy (Olympus Corporation, Tokyo, Japan) and the Metamorph imaging system (Molecular Devices, Downingtown, PA).

Immunofluorescence staining was performed using methods previously described [21]. CD31 and Collagen IV double staining were used to determine capillary. Seven μm cryostat sections were fixed in −20 °C methanol for 10 min. Sections were washed with PBS and blocked with 5 % normal goat serum before incubation at 4 °C with primary antibody. Secondary antibodies included goat anti-rabbit and donkey anti-rat IgG conjugated with Alexa Fluor 488 or Alexa Fluor 594 (Molecular Probes). Sections were examined using fluorescent microscope (BX41M-ESD, Olympus, Japan) and capillary density were quantified with the ImageJ software (National Institutes of Health, Bethesda, MD, USA).

### Western Blotting

Muscle tissue were homogenized and lysed with ice-cold buffer containing 25 mM Hepes (pH 7.5), 300 mM NaCl, 1.5 mM MgCl2, 0.2 mM EDTA, 0.1% Triton X-100, 20 mM β-glycerophosphate, 0.1 mM sodium orthovanadate, 0.5 mM DTT, 100 g/ml PMSF, and 2 g/ml leupeptin. Approximately 100 μg of protein was separated by electrophoresis in a 10% sodium-dodecyl-sulfate polyacrylamide gel. After electrophoresis, the protein was electro-transferred onto polyvinylidene fluoride membranes (Life Science, Boston, MA). Membranes were probed with antibodies specific to phospho-ERK, ERK, iNOS, e-NOS, phospho-eNOS, and HIF-1α. After probing with a horseradish peroxidase-conjugated secondary antibody, protein signals were visualized using enhanced chemiluminescence reagents (Amersham, Arlington Heights, IL). The calculation was achieved using a chemiluminescence imager (Fusion-SL 3500 WL), and the signals were quantified with the ImageJ software (National Institutes of Health, Bethesda, MD, USA).

### Statistical analysis

Results were expressed as mean ± standard deviation (S.D.). Statistical significance was evaluated by one-way analysis of variance (ANOVA). *p*< .05 was considered significant difference.

## Results

### Improvement of blood flow recovery and increased capillary density with LLLT

To examine the therapeutic effects of low level LASER on diabetic mice, we applied LLLT on HLI mouse model. After diabetes was induced, HLI model was performed and mice received low level LASER for five constitutive days. Three days after LASER therapy finished, the blood perfusion was measured by Laser Doppler Perfusion Imaging (LDPI). As shown in Fig 1, after 8 days, blood perfusion in affected side was 69±37% of unaffected side, and LLLT can enhance blood perfusion to 190±28%. In diabetic mice, perfusion image showed perfusion in affected side was 37±18% of contralateral side, and LLLT can enhance blood perfusion to 191±45%. LDPI revealed significantly enhanced blood perfusion in the groups received LLLT compared to control or diabetes alone groups (Fig. 1B).

**Fig 1.**
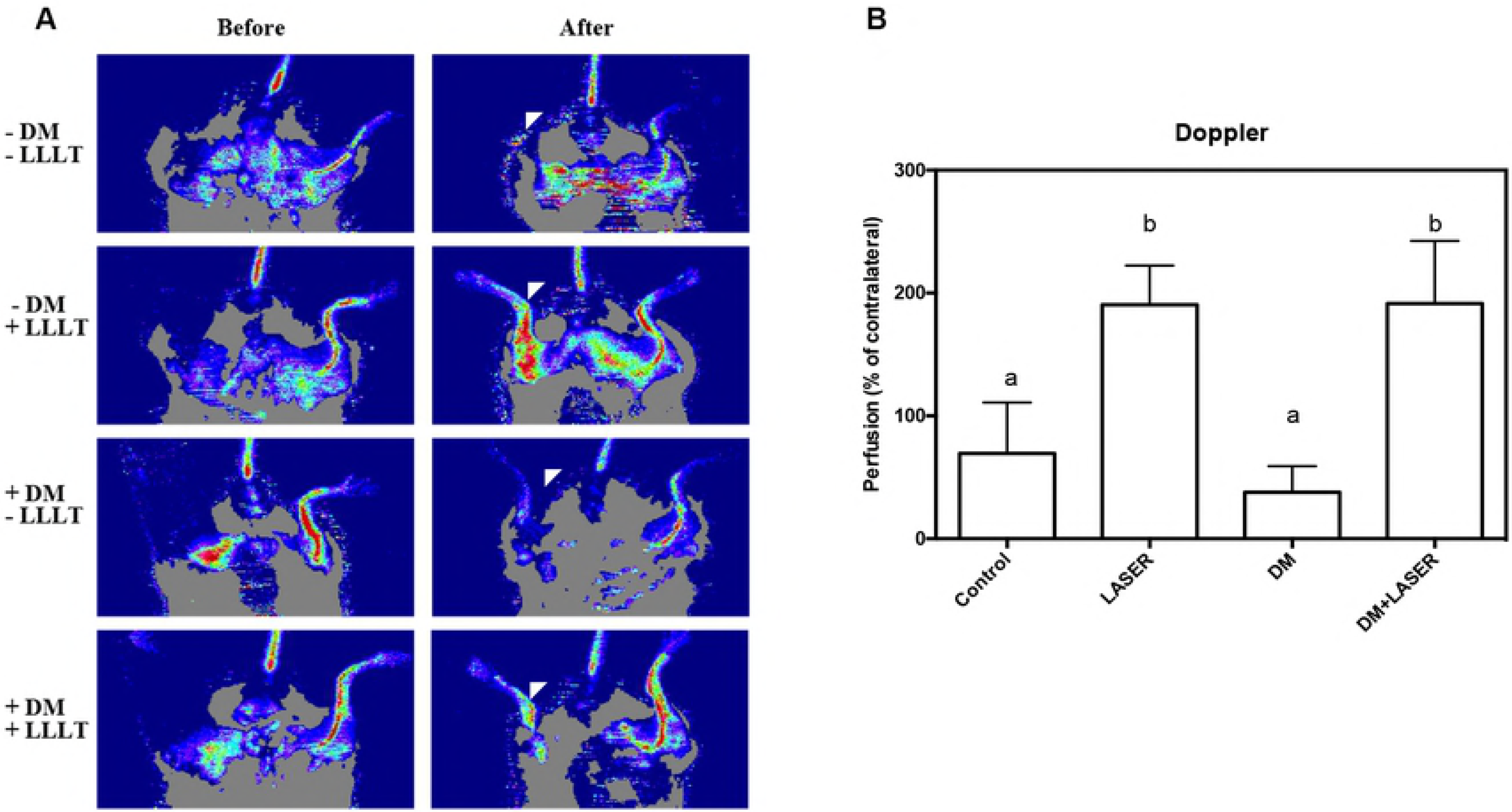
Effects of LLLT on perfusion recovery in HLI model on mice with or without diabetes. After diabetes and HLI were established, mice were treated with LLLT for constitutive five days. Representative LASER Doppler perfusion imaging images at 3 days later after LLLT finished (A); The perfusion ratio between the surgery and the control limb was calculated (B) Different symbols indicate statistically significant differences between treatments (*p* < 0.05).

We further measured capillary density to evaluate neovascularization. The hindlimb muscle were collected at 3 days after HLI and LLLT. The tissue was stained blood vessels with collage IV/CD31 and processed for fluorescence microscopic examination. Fig 2A shows the capillary density in the hindlimb muscle was significantly higher in the groups with LLLT (533±31/mm^2^) compared to those without (235±11.2/mm^2^). The capillary density was lower in mice with diabetes after HLI (157±29/mm^2^), and low level LASER notably increased capillary neovascularization in diabetic mice (410±60/mm^2^, Fig 2B).

**Fig 2.**
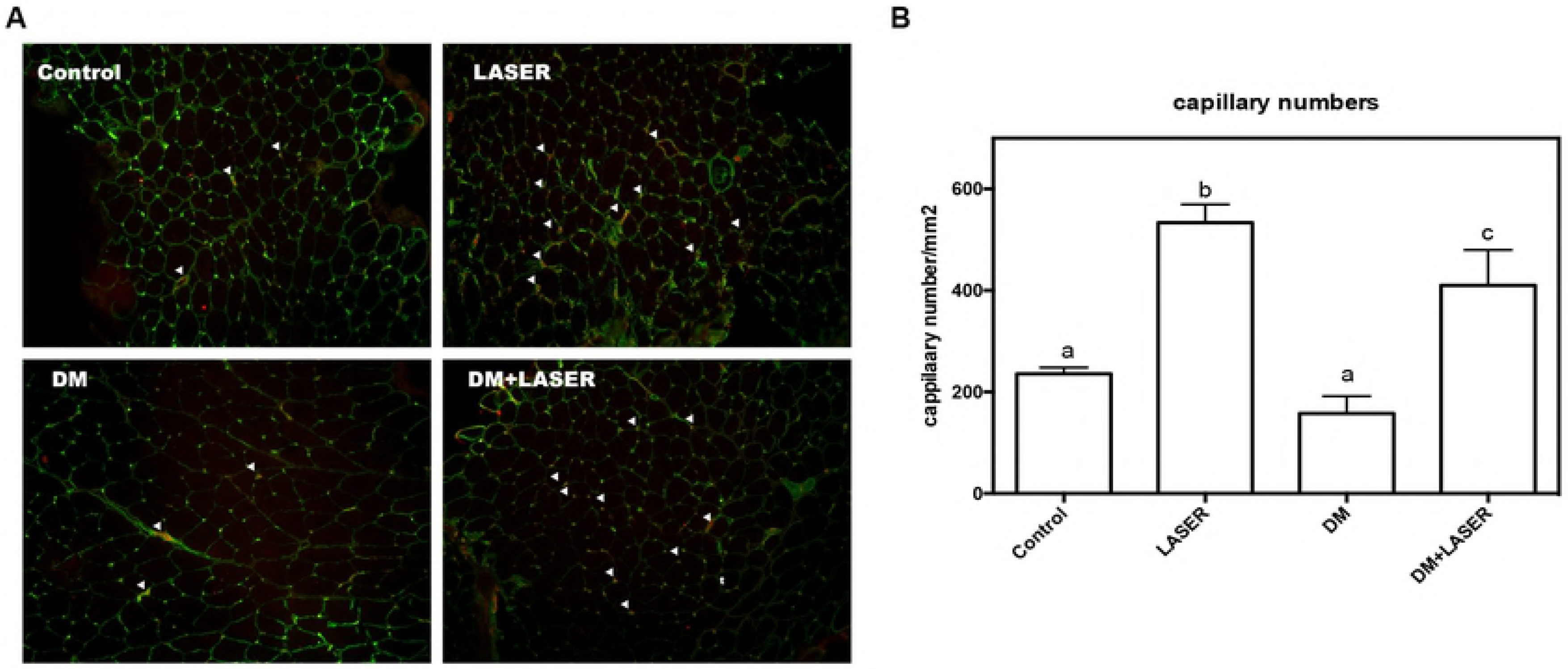
Effects of LLLT on capillary regeneration in in HLI model on mice with or without diabetes. After diabetes and HLI were established, mice were treated with LLLT for constitutive five days. Vascular structure was detected by immunofluorescence staining with anti-CD31 (red) and anti-Col IV (green) double stain. Arrow heads indicate capillaries (A); The data are expressed as the means ± the S.D. of five independent fields (B). Different symbols indicate statistically significant differences between treatments (*p* < 0.05).

### Elevated expression of CD31 and VEGF in mice receive LLLT

We next investigated the pro-angiogenic effects of low level LASER on ischemic hindlimb by determining expression levels of angiogenesis related proteins via Western blotting with muscle tissue at 3 days after LASER irradiation. CD31, also known as platelet endothelial cell adhesion molecule (PECAM-1), highly expresses on endothelial cells. In agree with a greater expression of CD31 measured by immunofluorescence staining, LLLT significantly increased CD31 protein level more than two-fold in muscles of mice no matter with or without diabetes.

Among angiogenic growth factors, vascular endothelial growth factor (VEGF) is the most well known one. VEGF expression promotes endothelial cell proliferation. Fig 3B shows in muscles with ischemia, the VEGF expression level is very low. After mice treated with low level LASER, the Western blot results show LASER massively increase VEGF protein expression compared to those without LLLT (15±0.6 fold of control level). In mice with DM, VEGF level induced by LLLT is slightly lower than mice without diabetes (11±2.2 fold of control level), but it is significantly higher than on the mice without LASER treatment (0.49±0.1 fold of control level).

**Fig 3.**
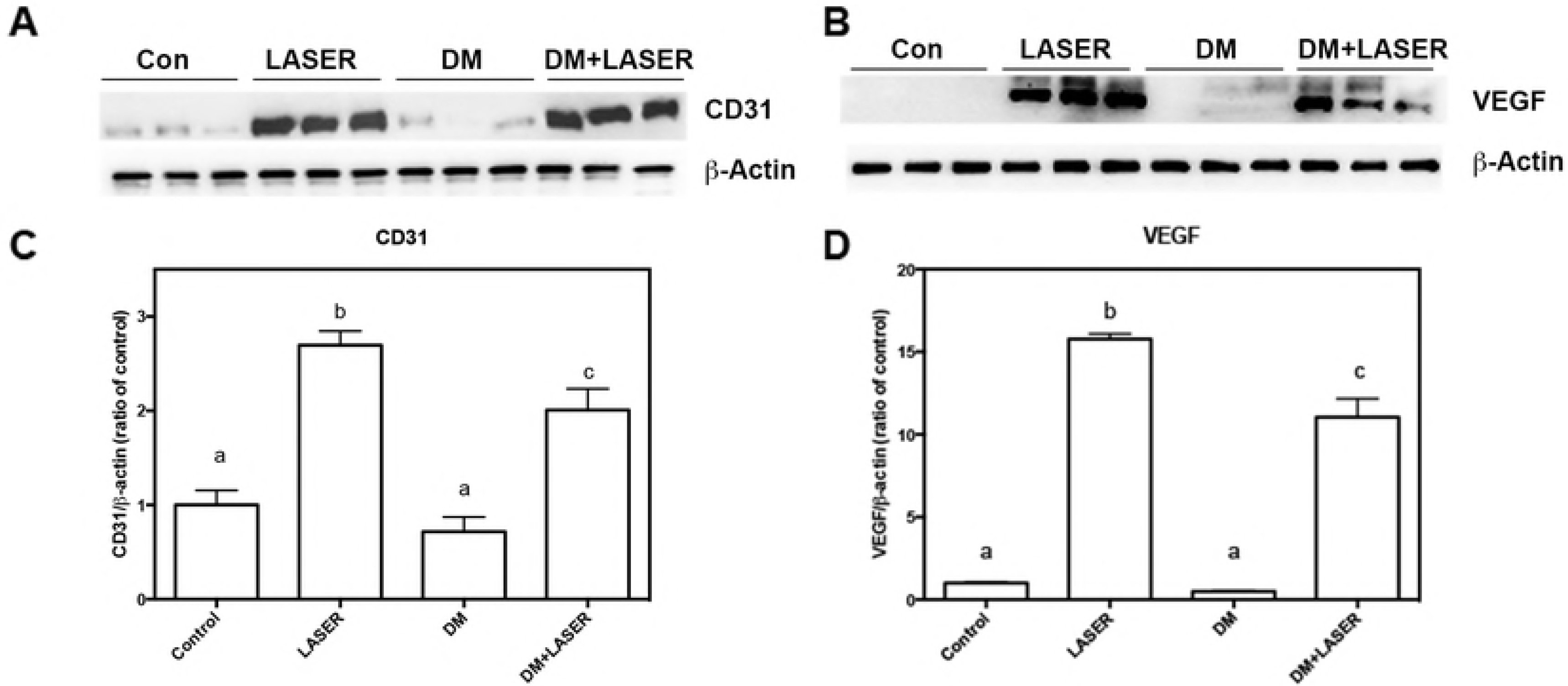
LLLT enhanced CD31 and VEGF in HLI mice with or without diabetes. After diabetes and HLI were established, mice were treated with LLLT for constitutive five days. Gastrocnemius/soleus muscles were harvested 3 days later. The levels of the CD31 (A) and VEGF (B) proteins were determined by Western blot; The quantifications of the related CD31 (C) and nVEGF (D) are shown. The values are presented as the means ± S.D. of three independent experiments. Different symbols indicate statistically significant differences between treatments (*p* < 0.05).

### The effects of LLLT on ERK MAPK signaling and HIF-1α in HLI mice model with diabetes

After VEGF binding to VEGF receptors, Raf/MEK/ERK pathway activates followed with HIF-1 α overexpression. These signal pathway plays key role in VEGF-mediated angiogenesis [22]. To confirm the effects of LLLT and its subsequent signalings, we further detected the activation of ERK and HIF-1α expression (Fig. 4). Western blot indicated that phospho-ERK was upregulated after 3 times LASER on mice no matter with or without diabetes, whereas the total amount of ERK and β-actin remained unchanged (Fig. 4A & 4B). As expected, HIF-1α protein levels were elevated in response to hypoxia in HLI control group compared to non-ischemic group (Fig. 4C & 4D). Similarly, HIF-1α protein expression was upregulated after LASER treatment both in LLLT and LLLT+DM groups.

**Fig 4.**
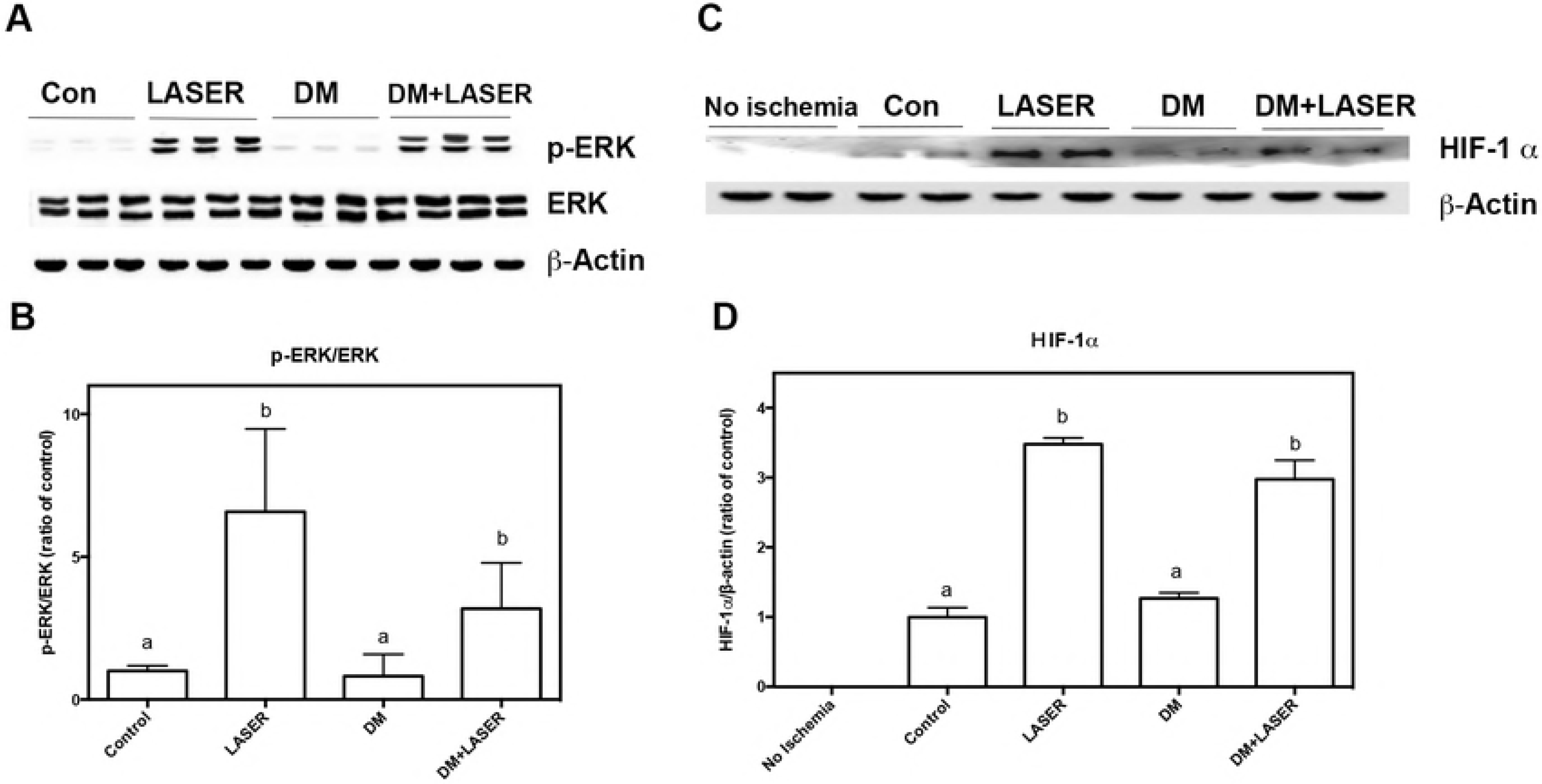
Effects of LLLT on ERK and HTF-1 α in HLI mice with or without diabetes. After diabetes and HLI were established, mice were treated with LLLT for constitutive five days. Gastrocnemius/soleus muscles were harvested 3 days later. The levels of the ERK and phospho-ERK (A) and HIF-1 α (B) were determined by Western blot; The quantifications of the related phospho-ERK/ERK ratio (C) and HIF-1 α (D) are shown. The values are presented as the means ± S.D. of three independent experiments. Different symbols indicate statistically significant differences between treatments (*p* < 0.05).

### Effect of LLLT on the Activation of eNOS and the Expression of iNOS in in HLI mice model with diabetes

We further investigated whether eNOS and iNOS pathways were involved in the LLLT induced angiogenesis. Fig 5A and 5B indicated that p-eNOS was upregulated after LASER treatment, whereas the total amount of eNOS remained unchanged (LLLT: 3±0.38 fold of control level). Previous studies show iNOS overexpression is associated with diabetes-induced endothelial dysfunction [23]. As expected, iNOS protein levels were significantly increased in response to diabetes induction (Figs. 5C and 5D), whereas low level LASER reduced these iNOS over-expression. Similarly, the expression level of iNOS was slightly lower in mice received LLLT (48±13% of control level) compared to those did not.

**Fig 5.**
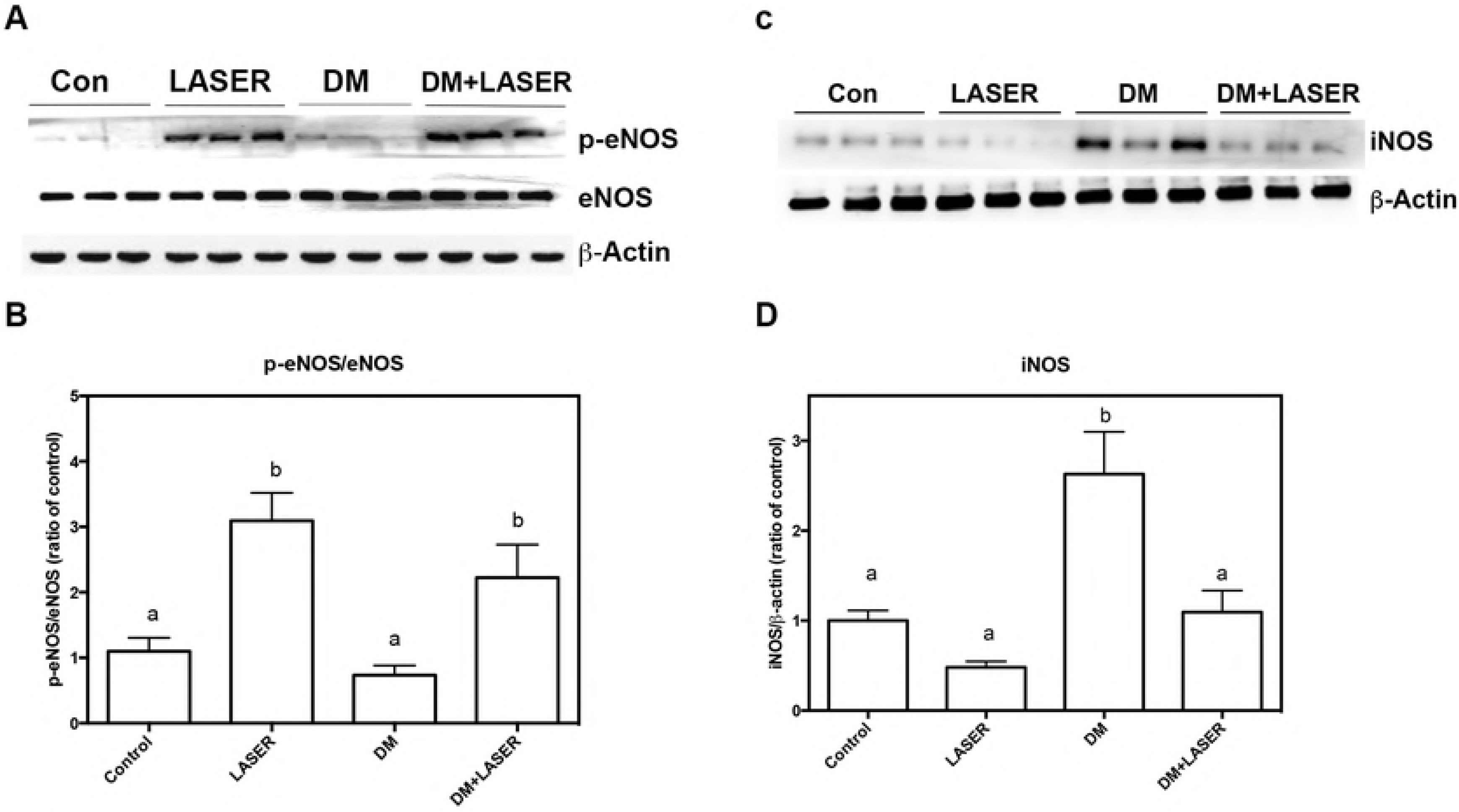
LLLT induced eNOS phosphorylation and decreased iNOS expression in HLI mice with diabetes. After diabetes and HLI were established, mice were treated with LLLT for constitutive five days. Gastrocnemius/soleus muscles were harvested 3 days later. The levels of the eNOS and phospho-eNOS (A) and iNOS (B) were determined by Western blot; The quantifications of the related phospho-eNOS/eNOS ratio (C) and iNOS (D) are shown. The values are presented as the means ± S.D. of three independent experiments. Different symbols indicate statistically significant differences between treatments (*p* < 0.05).

## Discussion

Here we have characterized the effect of low level LASER on angiogenic effects on diabetic mice with hind limb ischemia. Our findings demonstrate that LLLT promotes angiogenesis via increasing related signal molecules in the region of the muscles most severely affected in ischemia. The observed effects of low level LASER on peripheral artery occlusion may contribute to diabetes-related circulation issues.

Previous studies show low level LASER can improve skin flap viability through increasing angiogenesis rat ischemic skin flap model [17]. In addition to skin flap, here we demonstrated that LLLT could also promoting revascularization on diabetic mice with PAOD. Occlusion of femoral artery decreased the perfusion and capillary numbers of hindlimbs, which are more significant in mice with diabetes. In addition, ischemia plus high glucose decreased CD31 expression, which is the marker of vessels. LLLT reversed these harmful effects through enhancing VEGF expression and neovascularization.

Exposure to low level LASER significantly improved artery ligation-induced ischemia through promoting angiogenesis. The induction of pro-angiogenic factors play key roles in angiogenesis. Expression of VEGF is the major activator of angiogenesis in most condition. When VEGF is released, VEGF binds its receptor-2 (Flk-1/KDR) and strongly activates ERK 1/2 signals in different conditions [24]. These VEGF-ERK1/2 pathway plays crucial role in angiogenesis which demonstrated by is the results that endothelial tubulogenesis is disrupted by MEK inhibitor in vitro [25]. Both in adipose-derived mesenchymal stem cells and bone marrow derived stem cells, VEGF-induced stem cell differentiation into endothelial cell is controlled by ERK1/2 signaling [26] [27]. Blockage of ERK1/2 by specific inhibitor suppresses the expression of endothelial markers during differentiation. Through irradiation with LLLT, ERK phosphorylation and VEGF secretion are higher than non-treated human granulosa cells [28]. Thus, low level LASER stimulates VEGF and ERK endothelial cells and capillary regeneration, which are shown in different cell types and models.

In addition to VEGF activating ERK, ERK-activated HIF-1α signal also plays an important role[29]. As shown in Fig 4C, HLI alone slightly induced HIF-1α overexpression, and LLLT strongly activate HIF-1α as long along with ERK phosphorylation. Through ERK specific inhibitor, PD98059, evidences show activation and nucleocytoplasmic translocation of HIF-1α is controlled by ERK [30, 31]. ERK1/2 phosphorylates HIF-1α at S641 and S643, and these phosphorylated HIF-1 α are stabilized and translocate to nucleus [32, 33]. However, hypoxia upregulated HIF-1α transcription is independent to ERK1/2 MAPK in pulmonary artery smooth muscle cells [34]. Thus, we suggested that ERK 1/2 is in reposed to HIF-1 α activation in our model.

The balance between eNOS and iNOS takes parts in angiogenesis. The activation of eNOS plays a key role in endothelial cell function and survival. Nanomolar concentrations of NO produced by eNOS have anti-inflammatory and protective effects on endothelial cells [35]. Although NO can stimulate smooth muscle cells express VEGF both in human and rat, excessive NO produced by iNOS can cause endothelial cells apoptosis in ischemia/reperfusion injury [36, 37]. Our results demonstrated that LLLT exerts its protective effects by activating eNOS and strongly down-regulating iNOS expression which was dominant in diabetic mice, thereby suppressing the overproduction of NO.

In summary, we have demonstrated that LLLT promoting angiogenesis to against ischemia-induced apoptosis by promoting VEGF and eNOS signals. Our results provide therapeutic potential of LLLT on treating peripheral arterial diseases on diabetic patients.

## References

1. Ouriel K. Peripheral arterial disease. Lancet. 2001;358(9289):1257–64.

2. European Stroke O, Tendera M, Aboyans V, Bartelink ML, Baumgartner I, Clement D, et al. ESC Guidelines on the diagnosis and treatment of peripheral artery diseases: Document covering atherosclerotic disease of extracranial carotid and vertebral, mesenteric, renal, upper and lower extremity arteries: the Task Force on the Diagnosis and Treatment of Peripheral Artery Diseases of the European Society of Cardiology (ESC). Eur Heart J. 2011;32(22):2851–906.

3. Lord SR, Lloyd DG, Nirui M, Raymond J, Williams P, Stewart RA. The effect of exercise on gait patterns in older women: a randomized controlled trial. J Gerontol A Biol Sci Med Sci. 1996;51(2):M64–70.

4. Mohler ER, 3rd, Hiatt WR, Creager MA. Cholesterol reduction with atorvastatin improves walking distance in patients with peripheral arterial disease. Circulation. 2003;108(12):1481–6.

5. Antithrombotic Trialists C. Collaborative meta-analysis of randomised trials of antiplatelet therapy for prevention of death, myocardial infarction, and stroke in high risk patients. BMJ. 2002;324(7329):71–86.

6. Norgren L, Hiatt WR, Dormandy JA, Nehler MR, Harris KA, Fowkes FG, et al. Inter-Society Consensus for the Management of Peripheral Arterial Disease (TASC II). Eur J Vasc Endovasc Surg. 2007;33 Suppl 1:S1–75.

7. Tateishi-Yuyama E, Matsubara H, Murohara T, Ikeda U, Shintani S, Masaki H, et al. Therapeutic angiogenesis for patients with limb ischaemia by autologous transplantation of bone-marrow cells: a pilot study and a randomised controlled trial. Lancet. 2002;360(9331):427–35.

8. Folkman J. Seminars in Medicine of the Beth Israel Hospital, Boston. Clinical applications of research on angiogenesis. N Engl J Med. 1995;333(26):1757–63.

9. Lim W, Lee S, Kim I, Chung M, Kim M, Lim H, et al. The anti-inflammatory mechanism of 635 nm light-emitting-diode irradiation compared with existing COX inhibitors. Lasers Surg Med. 2007;39(7):614–21.

10. Bjordal JM, Johnson MI, Iversen V, Aimbire F, Lopes-Martins RA. Low-level laser therapy in acute pain: a systematic review of possible mechanisms of action and clinical effects in randomized placebo-controlled trials. Photomed Laser Surg. 2006;24(2):158–68.

11. Enwemeka CS, Parker JC, Dowdy DS, Harkness EE, Sanford LE, Woodruff LD. The efficacy of low-power lasers in tissue repair and pain control: a meta-analysis study. Photomed Laser Surg. 2004;22(4):323–9.

12. Hopkins JT, McLoda TA, Seegmiller JG, David Baxter G. Low-Level Laser Therapy Facilitates Superficial Wound Healing in Humans: A Triple-Blind, Sham-Controlled Study. J Athl Train. 2004;39(3):223–9.

13. de Loura Santana C, Silva Dde F, Deana AM, Prates RA, Souza AP, Gomes MT, et al. Tissue responses to postoperative laser therapy in diabetic rats submitted to excisional wounds. PLoS One. 2015;10(4):e0122042.

14. Loevschall H, Arenholt-Bindslev D. Effect of low level diode laser irradiation of human oral mucosa fibroblasts in vitro. Lasers Surg Med. 1994;14(4):347–54.

15. Lapchak PA, De Taboada L. Transcranial near infrared laser treatment (NILT) increases cortical adenosine-5′-triphosphate (ATP) content following embolic strokes in rabbits. Brain Res. 2010;1306:100–5.

16. Goralczyk K, Szymanska J, Lukowicz M, Drela E, Kotzbach R, Dubiel M, et al. Effect of LLLT on endothelial cells culture. Lasers Med Sci. 2015;30(1):273–8.

17. Cury V, Moretti AI, Assis L, Bossini P, Crusca Jde S, Neto CB, et al. Low level laser therapy increases angiogenesis in a model of ischemic skin flap in rats mediated by VEGF, HIF-1alpha and MMP-2. J Photochem Photobiol B. 2013;125:164–70.

18. Chu YH, Chen SY, Hsieh YL, Teng YH, Cheng YJ. Low-level laser therapy prevents endothelial cells from TNF-αlpha/cycloheximide-induced apoptosis. Lasers Med Sci. 2018;33(2):279–86.

19. Dai Q, Huang J, Klitzman B, Dong C, Goldschmidt-Clermont PJ, March KL, et al. Engineered zinc finger-activating vascular endothelial growth factor transcription factor plasmid DNA induces therapeutic angiogenesis in rabbits with hindlimb ischemia. Circulation. 2004;110(16):2467–75.

20. Shevchenko EK, Makarevich PI, Tsokolaeva ZI, Boldyreva MA, Sysoeva VY, Tkachuk VA, et al. Transplantation of modified human adipose derived stromal cells expressing VEGF165 results in more efficient angiogenic response in ischemic skeletal muscle. J Transl Med. 2013;11:138.

21. Enzmann G, Mysiorek C, Gorina R, Cheng YJ, Ghavampour S, Hannocks MJ, et al. The neurovascular unit as a selective barrier to polymorphonuclear granulocyte (PMN) infiltration into the brain after ischemic injury. Acta Neuropathol. 2013;125(3):395–412.

22. Chen JA, Shi M, Li JQ, Qian CN. Angiogenesis: multiple masks in hepatocellular carcinoma and liver regeneration. Hepatol Int. 2010;4(3):537–47.

23. Nagareddy PR, Xia Z, McNeill JH, MacLeod KM. Increased expression of iNOS is associated with endothelial dysfunction and impaired pressor responsiveness in streptozotocin-induced diabetes. Am J Physiol Heart Circ Physiol. 2005;289(5):H2144–52.

24. Wu LW, Mayo LD, Dunbar JD, Kessler KM, Baerwald MR, Jaffe EA, et al. Utilization of distinct signaling pathways by receptors for vascular endothelial cell growth factor and other mitogens in the induction of endothelial cell proliferation. J Biol Chem. 2000;275(7):5096–103.

25. Ilan N, Mahooti S, Madri JA. Distinct signal transduction pathways are utilized during the tube formation and survival phases of in vitro angiogenesis. J Cell Sci. 1998;111 (Pt 24):3621–31.

26. Xu J, Liu X, Jiang Y, Chu L, Hao H, Liua Z, et al. MAPK/ERK signalling mediates VEGF-induced bone marrow stem cell differentiation into endothelial cell. J Cell Mol Med. 2008;12(6A):2395–406.

27. Almalki SG, Agrawal DK. ERK signaling is required for VEGF-A/VEGFR2-induced differentiation of porcine adipose-derived mesenchymal stem cells into endothelial cells. Stem Cell Res Ther. 2017;8(1):113.

28. Kawano Y, Utsunomiya-Kai Y, Kai K, Miyakawa I, Ohshiro T, Narahara H. The production of VEGF involving MAP kinase activation by low level laser therapy in human granulosa cells. Laser Ther. 2012;21(4):269–74.

29. Berra E, Milanini J, Richard DE, Le Gall M, Vinals F, Gothie E, et al. Signaling angiogenesis via p42/p44 MAP kinase and hypoxia. Biochem Pharmacol. 2000;60(8):1171–8.

30. Minet E, Arnould T, Michel G, Roland I, Mottet D, Raes M, et al. ERK activation upon hypoxia: involvement in HIF-1 activation. FEBS Lett. 2000;468(1):53–8.

31. Hur E, Chang KY, Lee E, Lee SK, Park H. Mitogen-activated protein kinase kinase inhibitor PD98059 blocks the trans-activation but not the stabilization or DNA binding ability of hypoxia-inducible factor-1alpha. Mol Pharmacol. 2001;59(5):1216–24.

32. Mylonis I, Chachami G, Samiotaki M, Panayotou G, Paraskeva E, Kalousi A, et al. Identification of MAPK phosphorylation sites and their role in the localization and activity of hypoxia-inducible factor-1alpha. J Biol Chem. 2006;281(44):33095–106.

33. Triantafyllou A, Liakos P, Tsakalof A, Georgatsou E, Simos G, Bonanou S. Cobalt induces hypoxia-inducible factor-1alpha (HIF-1αlpha) in HeLa cells by an iron-independent, but ROS-, PI-3K- and MAPK-dependent mechanism. Free Radic Res. 2006;40(8):847–56.

34. Belaiba RS, Bonello S, Zahringer C, Schmidt S, Hess J, Kietzmann T, et al. Hypoxia up-regulates hypoxia-inducible factor-1alpha transcription by involving phosphatidylinositol 3-kinase and nuclear factor kappaB in pulmonary artery smooth muscle cells. Mol Biol Cell. 2007;18(12):4691–7.

35. Peng HB, Libby P, Liao JK. Induction and stabilization of I kappa B alpha by nitric oxide mediates inhibition of NF-kappa B. J Biol Chem. 1995;270(23):14214–9.

36. Zhu T, Yao Q, Wang W, Yao H, Chao J. iNOS Induces Vascular Endothelial Cell Migration and Apoptosis Via Autophagy in Ischemia/Reperfusion Injury. Cell Physiol Biochem. 2016;38(4):1575–88.

37. Dulak J, Jozkowicz A, Dembinska-Kiec A, Guevara I, Zdzienicka A, Zmudzinska-Grochot D, et al. Nitric oxide induces the synthesis of vascular endothelial growth factor by rat vascular smooth muscle cells. Arterioscler Thromb Vasc Biol. 2000;20(3):659–66.

